# Massively Parallel Reporter Assays identify enhancer elements in Oesophageal Adenocarcinoma

**DOI:** 10.1101/2024.06.11.598412

**Authors:** SH Yang, I Ahmed, Y Li, CW Bleaney, AD Sharrocks

## Abstract

Cancer is a disease underpinned by aberrant gene expression. Enhancers are regulatory elements that play a major role in transcriptional control and changes in active enhancer function are likely critical in the pathogenesis of oesophageal adenocarcinoma (OAC). Here, we utilise STARR-seq to profile the genome-wide enhancer landscape in OAC and identify hundreds of high-confidence enhancer elements. These regions are enriched in enhancer-associated chromatin marks, are actively transcribed and exhibit high levels of associated gene activity in OAC cells. These characteristics are maintained in human patient samples, demonstrating their disease relevance. This relevance is further underlined by their responsiveness to oncogenic ERBB2 inhibition and increased activity compared to the pre-cancerous Barrett’s state. Mechanistically, these enhancers are linked to the core OAC transcriptional network and in particular KLF5 binding is associated with high level activity, providing further support for a role of this transcription factor in defining the OAC transcriptome. Our results therefore uncover a set of enhancer elements with physiological significance, that widen our understanding of the molecular alterations in OAC and point to mechanisms through which response to targeted therapy may occur.

## Introduction

Oesophageal adenocarcinoma (OAC) is a gastrointestinal cancer that is one of the leading global causes of cancer-associated deaths [Coleman et al. 2018]. A number a genome-wide DNA-sequencing studies have identified potential disease driver events, including amplifications of genes encoding receptor tyrosine kinases (RTKs) such as ERBB2 [Frankell et al, 2019; Ross-Innes et al, 2015; Stachler et al, 2015]. Despite this, there is still a paucity of information regarding the specific mechanisms by which disease occurs. We have previously demonstrated that changes to the accessible chromatin landscape are critical in both the onset of disease, and the response to targeted therapeutic intervention [Britton et al, 2017; Rogerson et al, 2019; Rogerson et al, 2020; Ogden et al, 2022]. These chromatin changes often occur in regions of the genome that have the potential to function as regulatory elements, such as enhancers, and may represent an underappreciated mechanism by which disease and therapy-resistance is facilitated. This makes the study of enhancer elements in OAC of potential clinical importance.

Enhancers are distal regulatory elements that affect the expression of their target genes through long-range interactions with their promoters [Andersson & Sandelin, 2020]. The identification of enhancers has largely focused on the use of correlative approaches, linking the presence of histone marks such as H3K27ac and H3K4me1, as well as chromatin accessibility, to enhancer potential [Creyghton et al, 2010; Heintzman et al, 2009]. Whilst these approaches have revolutionised the study of enhancer elements, they remain hampered by the inability to provide a robust readout of genuine enhancer activity. Indeed, there is a body of evidence to suggest that correlative enhancer marks have no bearing on enhancer activity itself [Sankar et al, 2022; Pengelly et al, 2013; Dorighi et al, 2013]. Low-throughput enhancer reporter assays, which link enhancer function to a reporter gene such as green fluorescent protein (GFP) or luciferase, remain the most commonly used method by which enhancer activity is assessed [Inoue & Ahituv, 2015]. However, these approaches generally interrogate a single enhancer at time, limiting their use in genome-wide enhancer identification. To this end, genome-wide screening such as STARR-seq is an important advance in the understanding of enhancer biology [Arnold et al, 2013]. STARR-seq is a form of massively parallel reporter assay (MPRA), where regions of DNA are inserted downstream of a promoter, and into the 3’-UTR of a reporter gene. Provided that the region of DNA functions as an enhancer, it will drive expression of the reporter transcript, and itself, which may be detected using RNA-seq [Arnold et. al 2013]. By generating genome-wide libraries of potential regulatory DNA regions, and inserting these into the STARR reporter, it is possible to interrogate global enhancer activity.

Our previous studies identified regions of the genome that change accessibility during the transition to OAC from the precursor condition Barrett’s oesophagus [Britton et al, 2017; Rogerson et al, 2019; Rogerson et al, 2020], but these provide only correlative evidence for enhancer activity based on one modality (chromatin accessibility). Here, we employed STARR-seq to expand on these studies and provide direct evidence of enhancer activity in OAC. By using genome-wide DNA libraries generated using a combination of ATAC-seq and CUT&Tag, we identified chromatin regions in OAC that function as active enhancer elements. Integration with other genome-wide datasets from both OAC cells and patient samples validated their designation as *bona fide* regulatory elements and their association with regulatory events in OAC patients. These findings therefore expand our current understanding of the gene regulatory mechanisms underlying OAC.

## Results

### Identification of enhancer-like regulatory regions using STARR-seq in OAC cells

To identify genomic regions with enhancer activity in OAC cells, we applied STARR-seq in the OAC cell line, OE19 which we have previously demonstrated to faithfully recapitulate OAC at the chromatin level [Britton et al, 2017; Rogerson et al, 2019; Rogerson et al, 2020]. This technique leverages the ability of putative enhancer elements to drive transcription [Arnold et al, 2013; Inoue & Ahituv, 2015]. Putative enhancer elements are placed into the 3’-UTR of a truncated GFP reporter under the control of a minimal promoter (Fig. 1A). If these elements function as enhancers, they may upregulate the production of the reporter upon introduction into cells. Genomic sequencing of these cells can determine the extent of the plasmid introduction into the cells, whilst RNA sequencing can determine the extent of reporter RNA production. As the enhancers are within the 3’-UTR, they serve as their own barcode in the generated transcripts. By calculating the ratio of RNA to plasmid insertion, it is possible to calculate relative enhancer strength [Arnold et al, 2013; Inoue & Ahituv, 2015].

**Figure 1.**
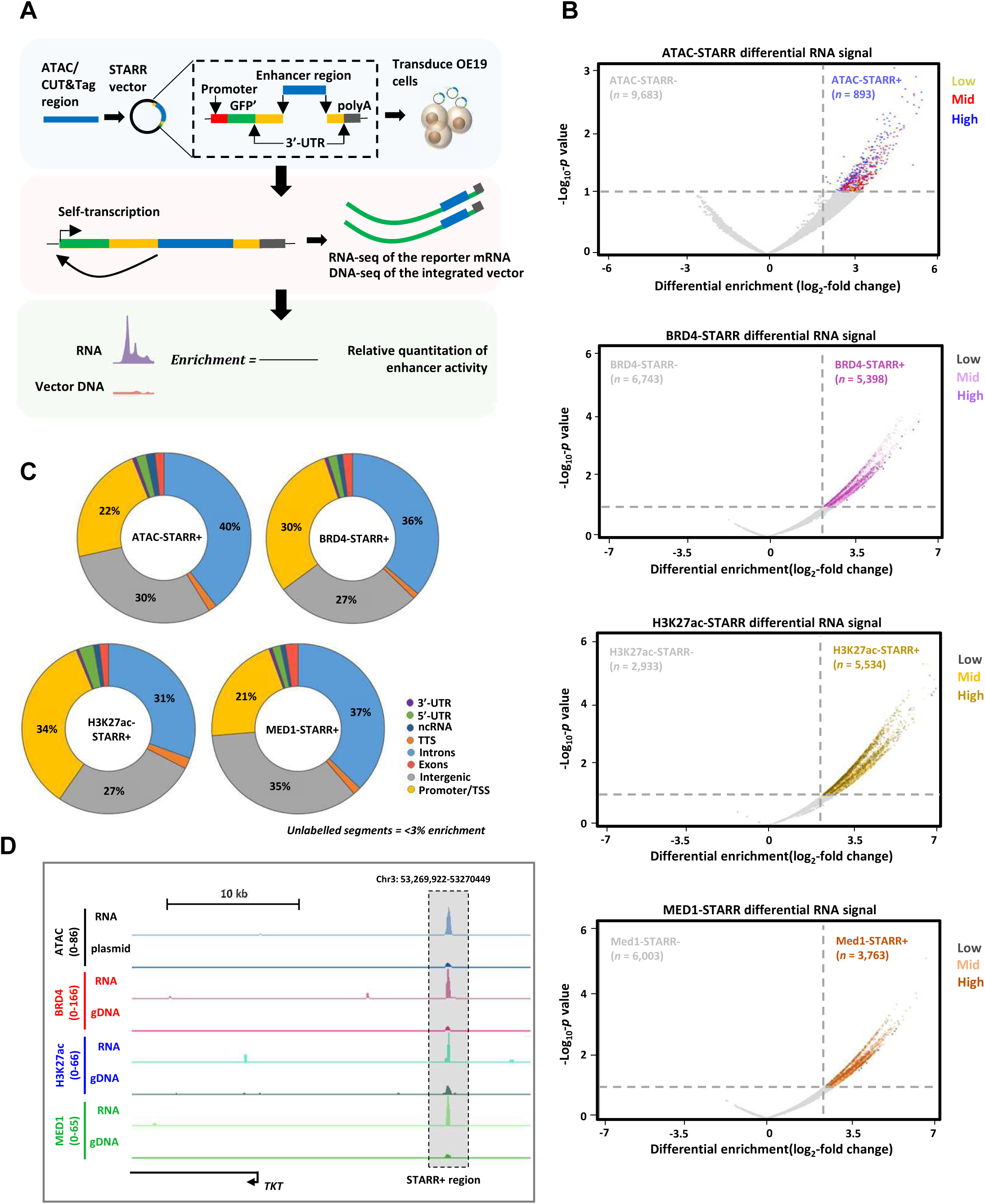
STARR-seq identifies potential enhancer regions. (A) STARR-seq strategy. Putative enhancer regions from ATAC-seq or CUT&Tag assays are inserted downstream from a truncated GFP reporter (GFP’) in a lentiviral vector. Regulatory regions which represent active enhancers are revealed by RNA-seq where they are “self transcribed” along with the GFP’ segment. Enrichment is calculated relative to input DNA from the library or integrated into the host genome. (B) Volcano plots displaying the differential (± Log_2_FC 2.0, < −Log_10_-*p*-value = 1.0) RNA signal over plasmid (ATAC) or gDNA (BRD4, H3K27ac, MED1) for ATAC, BRD4, H3K27ac and Med1-STARR-seq assays. Regions are colour coded by RNA signal strength. (C) Genomic distribution of the STARR+ regions for ATAC, BRD4, H3K27ac and MED1-STARR-seq assays. (D) Genome browser view of ATAC, BRD4, H3K27ac and MED1-STARR-seq assays, with plasmid (ATAC) or gDNA (BRD4, H3K27ac, MED1) and RNA tracks shown. Displayed is the *TKT* locus with an upstream STARR+ region highlighted.

We adopted a multi-pronged approach to identify putative enhancers by creating STARR-seq libraries from DNA fragments generated from OE19 cells, with accessible chromatin (via ATAC) or linked to the enhancer-associated chromatin factors and marks BRD4, H3K27ac and MED1 (via CUT&Tag). Libraries were created in lentiviral vectors to enable integration into the host genome. This created four independent libraries; ATAC-STARR, BRD4-STARR, H3K27ac-STARR and MED1-STARR libraries respectively. We then transduced OE19 cells with these libraries and used RNA-seq to identify enhancer derived transcripts alongside sequencing of the integrated gDNA as well as input plasmid library for determining signal enrichment relative to overall library abundance (Fig. 1A).

Differential STARR-derived RNA enrichment relative to plasmid library representation was calculated for each STARR-seq experiment and predominantly demonstrated higher levels of reporter-derived RNA signal enrichment, as expected (Fig. 1B). Genomic regions showing significant enrichment (log_2_≥2 fold; −log_10−_p-value ≤1) were taken forward as positive hits for demonstrating enhancer activity and henceforth be referred to as STARR+ regions (conversely those not exhibiting high signal are referred to as STARR-regions). This analysis revealed, 893, 5,398, 5,534 and 3,763 STARR+ regions representing potential enhancers, from the ATAC-STARR, BRD4-STARR, H3K27ac-STARR and MED1-STARR libraries respectively. These regions were predominantly from mid-high RNA producing regions rather than low activity, potentially high abundance regions as indicated by colour coding each region (Fig. 1B) based on whether in a low, mid or high expressing category (Supplementary Fig. S1A). Furthermore, we determined the impact of plasmid count on RNA production, by categorising STARR regions into low, mid and high plasmid DNA copy number regions (Supplementary Fig. 1B). This analysis demonstrated no disproportional enrichment of any plasmid DNA copy number categories, precluding plasmid drop-out or integration bias as drivers of RNA signal strength.

The genomic distribution of the STARR+ regions showed consistency across all STARR libraries (Fig. 1C) which is similar to that observed in all accessible ATAC-seq derived regions (Supplementary Fig. 1C). However, by comparison to input libraries (7-16%) an enrichment was observed for promoter elements (21-34%) (Fig. 1C). This phenomenon has previously been reported for other high throughput enhancer screens [Wang et al, 2018] and is consistent with the notion that a subset of promoters can function as enhancer elements in MPRAs [Andersson & Sandelin, 2020]. These observations confirm that the STARR+ elements are not unrepresentative elements predisposed to transduction, and are genuinely randomly sampled from the libraries from which they are generated.

STARR-seq therefore identifies a catalogue of self-transcribing regions that may represent enhancer elements in OAC cells. This is exemplied by the STARR+ region identified by all STARR-seq modalities, found at the *TKT* locus (Fig. 1D), which encodes a transketolase which has been associated with the tumourigenic properties of many different cancers [Hao et al, 2022].

### High-confidence STARR+ regions are marked with features of active chromatin

To create a high confidence set of STARR+ regions, we overlapped the regions found in each dataset (Fig.2A), and found 1,549 STARR+ intersect regions common to at least two of the ATAC, BRD4, H3K27ac and MED1 STARR+ datasets (Fig. 2A; Supplementary Table S1). These intersect STARR+ regions include promoter proximal elements in addition to both intergenic and intragenic regions (Supplementary Fig. S2A). Intragenic enhancers, in particular, can be challenging to identify through other approaches such as eRNA profiling in patient samples [Chen et al., 2018; Ahmed et al, 2023]. We next sought to identify the features associated with these STARR+ intersect regions, and the implications on regulatory potential. We previously generated CUT&Tag, ChIP-seq and ATAC-seq data for a range of chromatin-associated factors in OE19 cells [Ahmed et al, 2023; Rogerson et al., 2019; Rogerson et al., 2020] and we used this data to examine the overall levels of a broad range of chromatin marks and binding proteins across all of the STARR+ regions. Strikingly, the high confidence 1,549 STARR+ intersect peaks showed widespread enrichment for chromatin features associated with active regulatory potential, including BRD4, H3K27ac, MED1 and PolII as expected, but also more overt enhancer-associated marks such as H3K4me1/2 (Fig. 2B). This is exemplified by an intergenic region located upstream from *TKT* (Fig. 2C). Chromatin architectural marks were also detected (CTCF and SMC1) but there was little evidence of the repressive chromatin mark H3K27me3 (Fig. 2B). While the STARR+ regions identified in all of the individual assays exhibited enrichment for active chromatin features, the high confidence intersecting regions showed higher levels than any of the regions identified uniquely in a particular dataset (Fig. 2D; Supplementary Fig. S2B). This mirrors the data from the STARR-seq assays where there is high signal resulting from the assay in which the data is generated but high signal is only uniformly observed across all marks in the intersecting regions (Supplementary Fig. S2C and D). Overall, the high confidence intersect set of 1,549 STARR+ regions exhibit enrichment of features indicative of active enhancers in OE19 cells.

**Figure 2.**
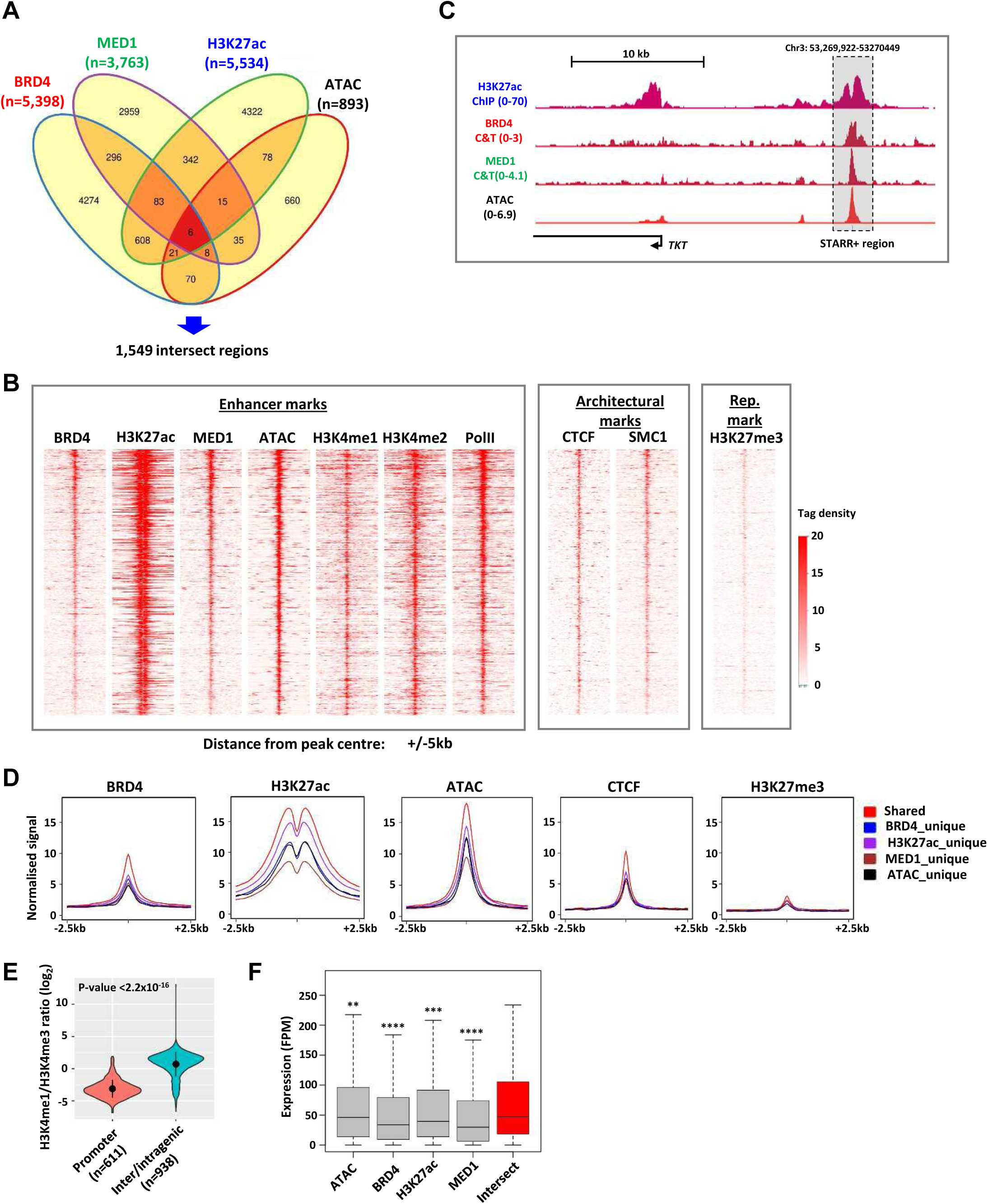
High confidence STARR+ regions are associated with active chromatin and transcriptional features. (A) Venn-diagram displaying the intersect between ATAC-, BRD4-, H3K27ac- and MED1-STARR+ regions. (B) Heatmaps showing CUT&Tag (BRD4, MED1, H3K4Me1, H3K4Me2, PolII, CTCF, SMC1 and H3K27me3), ChIP-seq (H3K27ac) and ATAC-seq signal in OE19 cells at the 1,549 STARR+ intersect regions. (C) Genome browser view of H3K27ac ChIP-seq, BRD4 CUT&Tag, MED1 CUT&Tag and ATAC-seq signal at an intergenic STARR+ intersect region highlighted upstream from the *TKT* locus. (D) Metaplots of CUT&Tag, ChIP-seq and ATAC-seq signal in OE19 cells at the 1,549 STARR+ intersect regions compared to STARR+ unique regions in ATAC, BRD4, H3K27ac and MED1-STARR-seq assays. (E) Violin plots displaying the ratio of H3K4me1:H3K4me3 ratios at distal regulatory and promoter proximal regions (−1kb to +0.1kb) within the 1,549 STARR+ intersect regions (*p*-value is shown; Welch’s t-test). (F) Box plots comparing the expression of genes in OAC patient tissue total RNA-seq samples from the OCCAMS dataset (n=210) annotated to unique BRD4, H3K27ac, MED1-STARR+ regions against genes annotated to the 1,549 STARR+ intersect regions (*p*-value is shown; **=<0.01, ***=<0.001, ****=<0.0001 Student’s t-test).

To further explore the characteristics of regions exhibiting enhancer activity, we compared the ratios of H3K4me1:H3K4me3 and H3K4me2:H3K4me3 found in OE19 cells between STARR+ intersect regions annotated as promoter-proximal (−1 kb to +0.1 kb) and the remaining intergenic or intragenic regions (representing distal enhancers). Both ratios were significantly higher for distal enhancer elements, than for promoter-proximal elements (Fig. 2E; Supplementary Fig. S2E). These analyses imply that the subset of intersect STARR+ regions located in intergenic or intragenic regions, may represent bona fide enhancers. This is corroborated by metaplots which show a clear demarcation of higher H3K4me1 signal at putative enhancers and H3K4me3 signal at promoter proximal regions (Supplementary Fig. S2F).

To explore the potential functional consequences of enhancer activity at the STARR+ intersect regions, we linked STARR+ regions to the nearest coding gene TSS using HOMER [Heinz et al, 2010] (Supplementary Table S1) and compared the expression of the 1,372 genes linked to the STARR+ intersect regions in OE19 cells and OAC patients, to those linked with the unique STARR+ regions. The intersect regions exhibited a significantly higher expression of associated genes compared to all of the other “unique” regions in both OE19 cells (Supplementary Fig. S2G) and OAC patients (Fig. 2F), consistent with a potential role as active enhancers for these genes in OAC.

In summary we have identified a high confidence dataset of regulatory elements in OAC cells with enhancer activity which are associated with high target gene activity in OAC patients.

### High-confidence STARR+ regions demonstrate enhancer activity

To further explore the enhancer-like properties of the STARR+ regions and more directly link the STARR+ regions to clinically relevant active transcription, we analysed kethoxal-assisted single-stranded DNA-sequencing (KAS-seq) data from OE19 OAC cells [Ahmed et al, 2023] and performed additional analyses on three OAC patient samples (Supplementary Table S2B). This approach serves as a proxy for active transcription, and potentially eRNA production, thereby indicative of enhancer activity given the close association between eRNA production and enhancer activity [Andersson & Sandelin, 2020]. The data from OE19 cells shows good correlation with each of the OAC tissue KAS-seq samples (Supplementary Fig. S3A). We focussed on intergenic regions to avoid interference from coding gene transcription and found substantially higher levels of KAS-seq signal in OE19 cells at the STARR+ intersect regions compared to a random set of accessible chromatin regions not identified in the STARR-seq assays (Supplementary Fig. S3B) consistent with enhancer transcription and hence enhancer activity in cancer cells. Next, we turned to patient samples, and again we found higher levels of KAS-seq signal associated with the intersect STARR+ regions, indicative of high enhancer activity in OAC patients (Fig. 3A). To further explore the relevance of the regulatory regions we have identified in a disease context, we next sought to determine whether any of our previously identified eRNAs in OAC patients [Ahmed et al, 2023] were located within the STARR+ intersect regions. Comparing the two datasets, we found a significant enrichment of eRNAs amongst the intergenic STARR+ intersect regions, consistent with potential enhancer activity in OAC patients (Fig. 3B).

**Figure 3.**
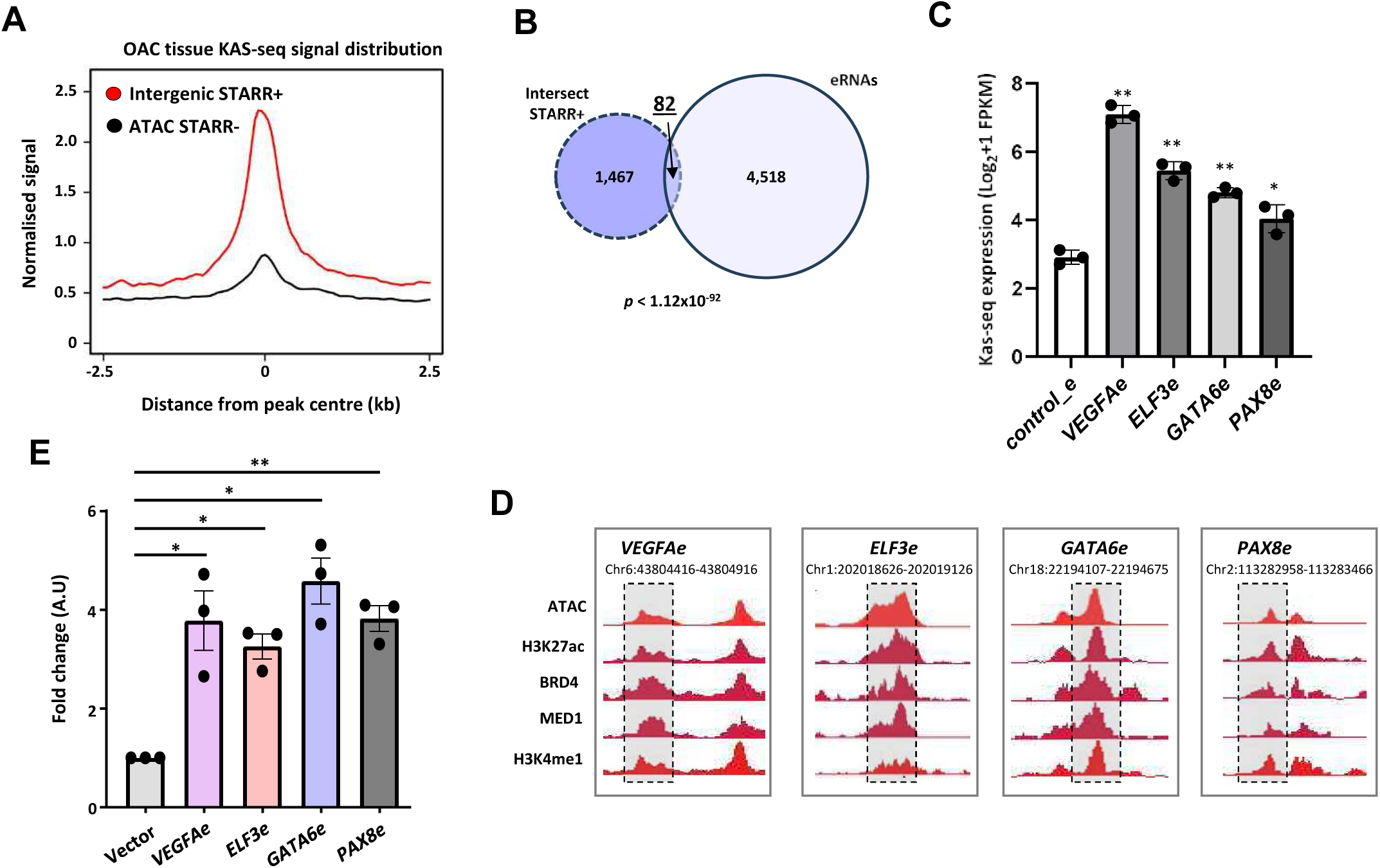
High confidence STARR+ regions are associated with enhancer-like activity. (A) Metaplot of OAC patient tissue KAS-seq signal at the 566 STARR+ intersect regions found in intergenic regions compared to 2,303 randomly selected STARR-ATAC peaks from OE19 cells. (B) Venn-diagram of overlap between the 1,549 STARR+ intersect regions and 4,600 previously identified BO and OAC patient eRNAs (*p-*value is shown; Fisher’s exact test). (C) Bar graph displaying the difference in KAS-seq expression between *VEGFAe, ELF3e*, *GATA6e* and *PAX8e* regions, compared to the average of 4 negative control regions in OE19 cell KAS-seq data (** = *p* < 0.01; * = *p* < 0.05; t-test relative to controls; n=3). (D) Genome browser view of ATAC-seq, BRD4, H3K27ac and MED1 and H3K4me1 CUT&tag signal at the regions associated with the *VEGFAe, ELF3e*, *GATA6e* and *PAX8e* STARR+ regions (boxed). (E) Bar graph displaying the difference in luciferase reporter activity between *VEGFAe, ELF3e*, *GATA6e* and *PAX8e,* compared to vector only negative control (** = *p* < 0.01; * = *p* < 0.05; one-way ANOVA with Bonferroni’s correction).

Next, we wished to validate a panel of enhancer regions from the STARR+ regions using an independent reporter assay based on a luciferase reporter in OE19 cells. We selected four STARR+ intersect regions, annotated to cancer relevant genes: *vascular endothelial growth factor A* (*VEGFA)*, *ELF3*, *GATA6* and *paired box gene 8* (*PAX8)*, which displayed significantly increased KAS-seq signal in OAC tissue relative to a panel of control genomic regions (Fig. 3C). These regions all show a range of chromatin features indicative of enhancer activity (Fig. 3D). Importantly, all regions demonstrated a significant increase in luminescence in the reporter assay, relative to vector only control (Fig 2E) authentic enhancer activity.

Collectively, this data further supports the conclusion that the STARR+ intersect regions we have identified represent *bona fide* regulatory elements with enhancer-like properties and demonstrates their likely clinical relevance due to their association with patient-derived eRNAs and KAS-seq signal in OAC tissue.

### Transcription factor and gene activity associated with high-confidence STARR+ regions in OAC cells and patients

To further understand the biological activity associated with the high confidence STARR+ regions, we performed gene ontology (GO) analysis on their associated genes. This revealed a significant enrichment in processes associated with OAC, including “VEGFA signalling”, “pathways in cancer” and “actin cytoskeleton organisation” (Fig. 4A). Next, we sought to identify the transcription factors that may bind and regulate the STARR+ intersect regions. We identified an enrichment of DNA motifs associated with transcription factors previously shown to play a role in OAC, such as KLF5 as well as AP-1, FOX and ETS family members (Fig. 4B) [Rogerson et al., 2020; Britton et al., 2017; Rogerson et al., 2020; Chen et al., 2020]. In the case of KLF5, motif occurrences in the STARR+ regions were significantly higher than across all open chromatin regions in OE19 cells, whereas other transcription factors previously implicated in OAC either showed no enrichment (HNF4A) or reduced enrichment (GATA6) relative to the expected frequency (Fig. 4C). To provide more direct evidence for regulatory potential, we examined overlaps between ChIP-seq binding datasets for KLF5, GATA6 and HNF4A transcription factors in OE19 cells at the peak (Supplementary Fig. S4A) or signal level (Fig. 4D). In all cases, significant overlap was observed (Supplementary Fig. S4A). Subsequent clustering of binding activity allowed us to identify five broad subgroups among the STARR+ regions (Fig. 4D), with a relatively small triply bound group (C1:KGH), a group co-bound bound by KLF5 and GATA6 (C2:KG), two groups bound by either KLF5 (C3:K) or GATA6 alone (C4:G) and one group not bound at high levels by any of the transcription factors (C:5). Regions surrounding the *VEGFA* (Fig. 4E) and *PIM3* (Supplementary Fig. S4B) gene loci exemplify STARR+ regions from the C1:KGH and C3:K clusters respectively. All groups showed high levels of open chromatin, H3K27ac and BRD4 binding (Fig. 4D, right; Supplementary Fig. S4C, top). However, KAS-seq signal (an indicator of active transcription) was highest in the KLF5 only cluster (K) and lowest in the cluster featuring GATA6 binding (G) (Fig. 4D, right; Supplementary Fig. S4C, bottom left). Importantly, this high signal in the KLF5-specific cluster was maintained when considering only intergenic STARR+ regions (Supplementary Fig. S4C, bottom right) indicating that this signal is likely generated from eRNA transcription, a feature of active enhancers (Andersson et al., 2014). We therefore analysed KAS-seq from OAC patient samples and found that the KLF5-specific STARR+ regions also exhibited the highest transcriptional activity, consistent with active enhancer activity in patients (Fig. 4F).

**Figure 4.**
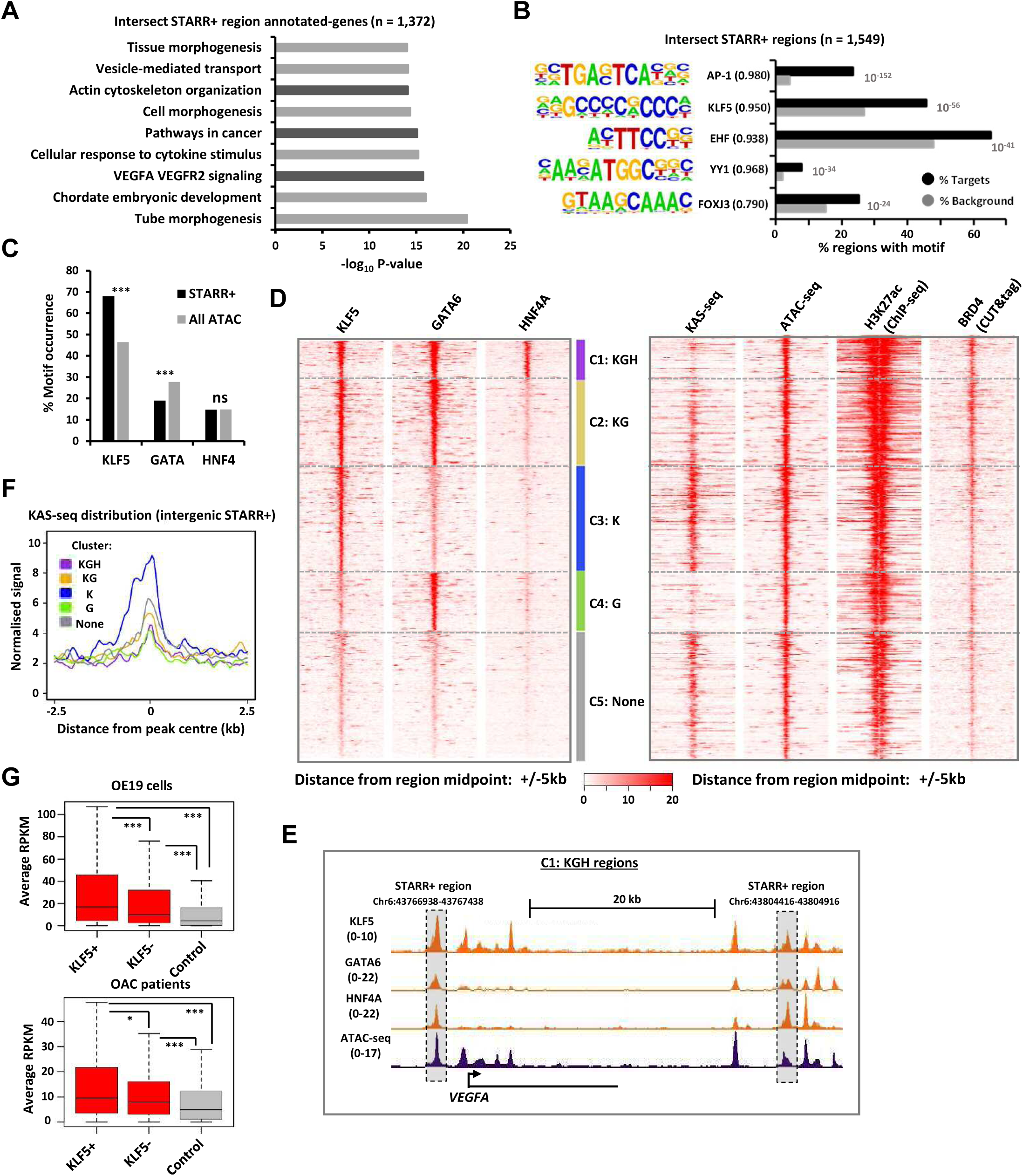
High confidence STARR+ regions are enriched for binding to core OAC transcription factors, active chromatin features and are associated with active genes. (A) GO-term analysis of genes annotated to the 1,549 STARR+ intersect regions. (B) Transcription factor *de novo* motif enrichment using HOMER for STARR+ intersect regions (*p*-values are shown). Matches to the indicated motifs are indicated in brackets. (C) Frequency of DNA binding motif occurrence for KLF5 (DGGGYGKGGC), GATA (NBWGATAAGR) and HNF4 (CARRGKBCAAAGTYCA) transcription factors in the 1,549 intersect STARR+ regions and all 99,855 ATAC-seq peaks found in OE19 cells. **** = P-value<0.001; ns = non-significant. (D) Heatmap of ChIP-seq signal for the transcription factors KLF5 (K), GATA6 (G) and HNF4A (H) after kmeans clustering into 5 clusters (left) and associated KAS-seq, ATAC-seq, ChIP-seq and CUT&Tag signals for the same regions (right). (E) Genome browser view of KLF5, GATA6 and HNF4A ChIP-seq, and ATAC-seq signal at C1:KGH cluster STARR+ intersect regions (highlighted) surrounding the *VEGFA* locus. (F) Average tag density plots of KAS-seq signal from OAC patients (n=3) for each of the 5 clusters of the intergenic high confidence STARR+ regions. (G) Boxplot of average expression values of the genes nearest to the 1,549 STARR+ intersect regions from clusters C1-3 combined (KLF5+; 860 peaks, 774 genes), clusters C4-5 combined (KLF5-; 689 peaks, 659 genes) or 3 control sets of randomly selected ATAC-seq peaks, in OE19 cells (top) or 28 OAC patient samples (bottom). Statistical significance is shown; P-values <0.001, **** and <0.05, *.

Both motif analysis and ChIP-seq therefore implicate KLF5-associated STARR+ regions as active enhancers in OAC cell lines and patients. We therefore asked whether likely target genes also showed enhanced expression in OE19 cells. Compared to a random selection of open chromatin regions, all transcription factor binding-defined clusters showed significantly higher expression of their nearest genes, with the KLF5-specific cluster the highest level and the GATA6-specific cluster the lowest (Supplementary Fig. S4D, top). More generally, the genes closest to the KLF5-associated STARR+ regions showed higher expression in OE19 cells than those with no associated KLF5 binding (Fig. 4G, top). Similarly, in OAC patient samples a similar trend was observed with genes closest to KLF5-associated regions showing the highest expression (Fig. 4G, bottom; Supplementary Fig. S4D, bottom) but genes associated with GATA6-only regions showed the lowest expression levels in patients (Supplementary Fig. S4D, bottom).

Together these analyses demonstrate that STARR+ intersect regions are associated with OAC-specific transcription factors and have target genes that are related to OAC pathology. In particular, the association of the most active enhancers and their likely target genes with KLF5 binding is consistent with our previous finding of an important role for KLF5 in OAC progression from the Barrett’s pre-cancer state (Rogerson et al., 2019).

### High-confidence STARR+ regions are associated with clinically relevant regulatory events

The locus encoding the RTK ERRB2 is frequently amplified in OAC, and is therefore thought to be an oncogenic driver event [The cancer genome atlas research network, 2017; Frankell et al., 2019] and is a target for pharmacological inhibition in the clinic (Bang et al., 2010). We therefore explored whether the activity of the STARR+ regions was altered by treatment with the ERBB2 small-molecule inhibitor lapatinib. First, we assessed differential accessibility through ATAC-seq and H3K27ac signal (an activation associated histone mark) by ChIP-seq [Ogden et al, 2022] at STARR+ intersect regions upon lapatinib treatment for 24 hrs. Through using these datasets, we identified 2 and 24 STARR+ intersect regions that gained and lost accessibility, respectively, upon lapatinib treatment and 5 and 48 STARR+ intersect regions that increased and decreased in H3K27ac signal upon lapatinib treatment (Fig. 5A; Supplementary Table S3). Generally consistent changes were seen in chromatin accessibility and histone H3K27 acetylation levels, with decreases in both observed following lapatinib treatment (Fig. 5B). An exemplar region is located upstream from the *DUSP5* locus, where chromatin changes indicative of enhancer inactivation (Fig. 5C) accompanies reductions in gene expression (Supplementary Fig. S5A) following lapatinib treatment. Expression of *DUSP5* has potential clinical and prognostic significance as it is expressed more in OAC compared to Barrett’s (Supplementary Fig. S5B), and higher expression levels predict poorer patient survival (Fig. 5D).

**Figure 5.**
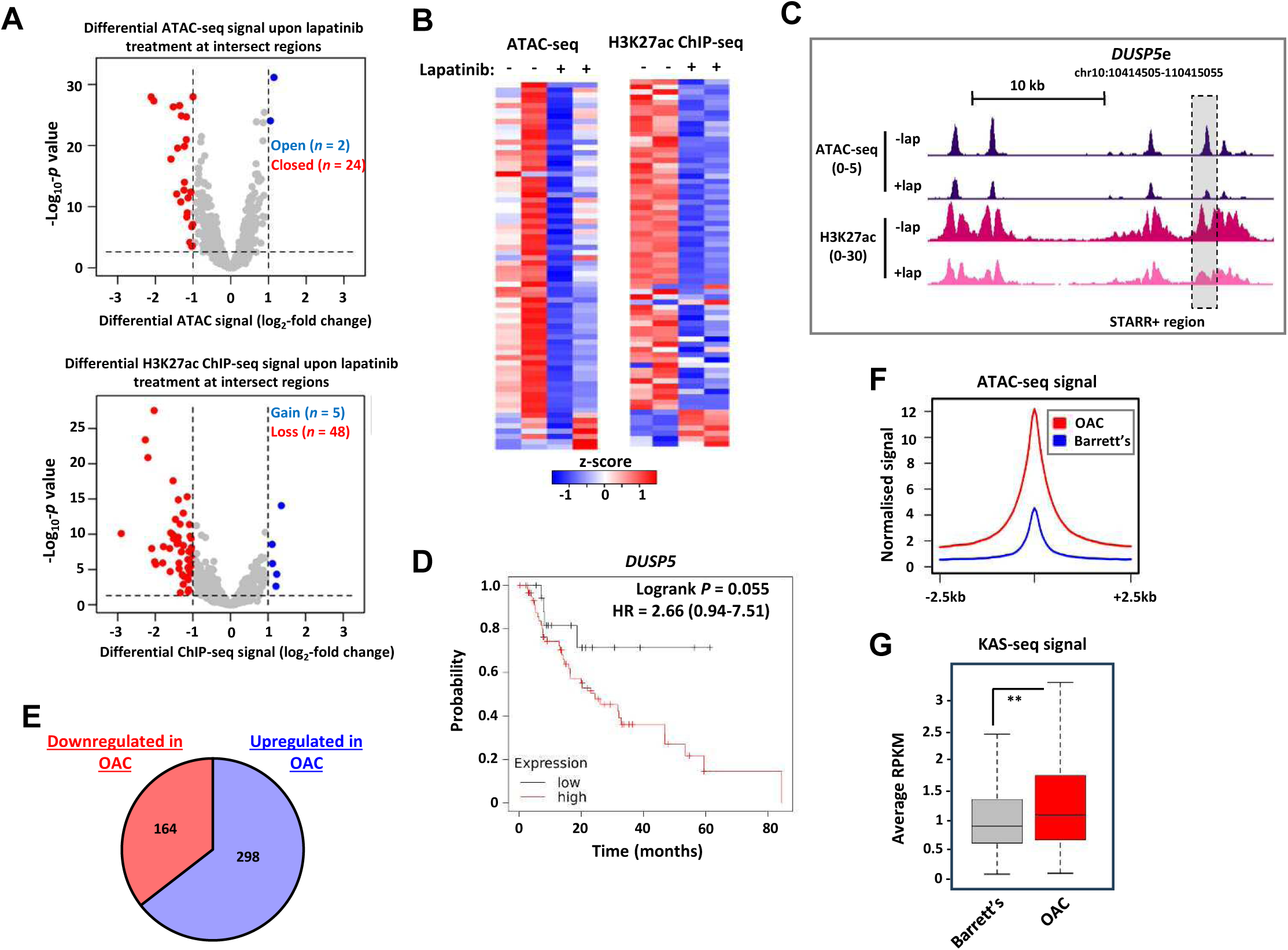
STARR+ defined enhancer regions respond to ERBB2 inhibition and are associated with OAC-specific gene regulation. (A) Volcano plots displaying the differential OE19 cell ATAC-seq (left; ± Log_2_FC=1, <*p-*value = 0.05) or OE19 cell H3K27ac ChIP-seq (right; ± Log_2_FC=1, < *p-*value = 0.5) at the 1,549 STARR+ intersect regions upon 24hr lapatinib treatment. (B) Heatmap of the ATAC-seq (left) or H3K27ac ChIP-seq (right) signal following 24hr lapatinib treatment of OE19 cells at the union of significantly closed/lost peaks from part A. Data are row z-normalised, with ATAC-seq and ChIP-seq data normalised separately. (C) Genome browser view of ATAC-seq and H3K27ac CUT&Tag signals at the STARR+ region upstream from *DUSP5* (*DUSP5*e) in OE19 cells following treatment with DMSO or lapatinib for 24hr. (D) Kaplan-Meier plot comparing overall survival between OAC patients (n=80) with low (n=19) or high (n=61) *DUSP5* expression in the TCGA PanCancer Atlas dataset (Log rank *p*-value is shown). (E) Pie chart showing the numbers of genes associated with the high confidence 1,549 STARR+ regions that are up or downregulated in OAC compared to matched Barrett’s samples from the same patients (n=28; >1.5 fold change, P-value<0.01). (F) Metaplots of ATAC-seq signal within the 1,549 STARR+ intersect regions in 4 Barrett’s or 11 OAC patient samples. (G) Boxplot of average KAS-seq signal in the 309 STARR+ intersect regions found in intergenic regions, from three matched Barrett’s and OAC patient samples. Statistical significance is shown; P-values <0.01, **.

Finally, we asked more globally whether the STARR+ regulatory regions we have identified are relevant to OAC by first asking whether accessible chromatin corresponding to these regulatory regions is also present in OAC patient samples. Almost all regions (98%) intersect with accessible chromatin peaks in at least one OAC patient and the majority are found in multiple patients, with 899 (58%) found in 8 or more patients (Supplementary Fig. S5C). Next, we compared the expression of their closest genes in both Barrett’s and OAC patients (Supplementary Table S2A) and there was clear enrichment of genes which are upregulated in OAC patients compared to the pre-cancer state (Fig. 5E). Furthermore, consistent with this observation, STARR+ regions are more accessible (Fig. 5F) and KAS-seq signal is higher at intergenic STARR+ regions in OAC patients compared to Barrett’s patients (Fig. 5G; Supplementary Table S2B), indicating higher enhancer activity in cancer cells

Collectively, this data demonstrates that the STARR+ regulatory regions we have identified are associated with OAC-specific properties in patients, as many are activated in response to oncogenic ERBB2 signaling and their activity is elevated in OAC patients. Our findings therefore provide us with insights into the active regulatory regions of the genome associated with this disease.

## Discussion

Cancer can be viewed as a disease characterised by disrupted gene regulatory networks [Zhao et al., 2021]. Enhancer elements represent fundamental hubs within these gene regulatory networks. When active, these regions serve as platforms for the integration of signals that dictate a broad range of outcomes influencing cellular phenotype, and when abnormally activated can result in tumourigenesis. Traditionally, histone marks such as H3K27ac and H3K4me1, in addition to chromatin accessibility, have been used to define enhancer elements [Creyghton et al, 2010; Heintzman et al, 2007]. However, while valuable, this approach is correlative; histone marks and accessibility may outline the location of a putative enhancer but these strategies remain limited for the definition of *bona fide* active enhancers. More recently, the finding that when enhancers are active they are themselves transcribed to produce eRNAs, has opened up the possibility of using eRNA profiling to identify enhancers in cancer cells (Chen et al., 2018; Ahmed et al, 2023). However, while promising, Intragenic enhancers can be challenging to identify through this approach due to the interference from ongoing genic transcription. Here, we used STARR-seq to identify representing potentially active enhancers in the OE19 OAC cell line. Using KAS-seq as a validation tool, in addition to our previously published patient eRNA dataset, we confirmed these regions as high confidence enhancers that are also operational in OAC patients. Moreover, we demonstrate the response of a subset of identified enhancers to the ERBB2 inhibitor lapatinib, which highlights the opportunity to approach clinically-relevant questions using our enhancer identification strategy.

A recent study applied STARR-seq to interrogate enhancer activity in gastric cancers [Sheng et al, 2021]. Whilst OAC does bear molecular similarities to a specific subtype of chromosomally-instable (CIN) gastric cancers [TCGA, 2017], the authors utilise a heterogeneous panel of gastric tumours and normal biopsies, as well as 28 gastric cancer cell lines to generate a STARR reporter library consisting of universal-common H3K27ac-marked elements. This amalgamation of sample types is likely to underestimate any findings given the exquisite cell-type specificity of enhancer activity. Additionally, it is now widely accepted that H3K27ac alone represents a poor indicator of enhancer activity [Pengelly et al, 2013; Sankar et al, 2022]. In addition to the lack of CIN mutational signature designation for the library source, these caveats preclude the general applicability of these findings to OAC. Here, we focused on using OE19 cells, which faithfully recapitulate OAC at the chromatin level [Britton et al, 2017; Rogerson et al, 2019; Rogerson et al, 2020]. Furthermore, our strategy for STARR library generation uses a range of input material that are associated with enhancers, and includes H3K27ac but also BRD4, MED1 and accessible chromatin. Collectively, this approach ensures the relevancy of our results, and broadens the search radius, improving chances of successfully identifying active enhancer regions.

Our approach identified ~1,500 high confidence active enhancers of potential relevance to OAC. These enhancer elements displayed an association with genes linked to processes pertinent to OAC biology, as well as an enrichment in the motifs of transcription factors which have been shown to play a role in the development of OAC [Rogerson et al., 2020; Britton et al., 2017; Rogerson et al., 2020; Chen et al., 2020]. Integration with our previously published genome-wide binding data on a variety of chromatin-associated factors in OE19 cells, as well as our patient-derived eRNA dataset, we verified the enhancer-like nature of our ~500 regions in addition to highlighting their potential clinical relevance. In support of this, we generated KAS-seq data from OAC patient tumour samples to monitor enhancer activity. KAS-seq measures the transcription bubble [Wu et al, 2020], serving as a proxy for enhancer transcription. Accordingly, these regions demonstrated high KAS-seq signal in both OE19 cells and patients further confirming their designation as high confidence enhancers. However, we also identified a large number of putative enhancers that are unique to a particular STARR assay. Further data mining alongside newly generated datasets will likely uncover additional *bona fide* enhancers in these cohorts, providing more broader utility of this resource to the community.

An interesting observation from our data was that KLF5 motif presence and chromatin binding is associated with more active enhancer regions and associated target genes whereas GATA6 bound enhancers exhibit the opposite tendency. The former observation is consistent with our previous finding that KLF5 repurposing through redistribution to a novel set of regulatory regions is one of the key molecular events that distinguishes OAC from the Barrett’s precursor state (Rogerson et al., 2020). Conversely, the implications on GATA6 functionality are less obvious as the gene encoding GATA6 is often amplified during the same transition, so a more prominent role in driving enhancer activity would be expected. It is possible that it acts to moderate the outputs of other transcription factors in the core OAC network to provide the optimal level of gene expression and this enigmatic phenomenon warrants further investigation alongside the outcomes of other combinatorial transcription factor interactions. Our dataset provides candidate regions to test these models.

To explore any clinical associations with our new STARR-seq-defined dataset, we utilised the small-molecule ERBB2 inhibitor, lapatinib. We have previously demonstrated that OE19 cells undergo genome wide chromatin accessibility changes upon lapatinib treatment, and this has important consequences for development of therapeutic resistance [Ogden et al, 2022]. These chromatin changes may reflect the activation or inactivation of enhancer elements. By integrating H3K27ac ChIP-seq and ATAC-seq data, we identified a subset of our newly defined enhancer elements that lose accessibility and become less marked by H3K27ac following lapatinib treatment. Furthermore, we show clear associations with increased accessibility and activity in OAC relative to Barrett’s patients, further emphasising the relevance of our data to gene regulatory events occurring in OAC patients.

In summary, our STARR-based approach has allowed us to identify a set of enhancers that are active in OAC which has led to a wider understanding of the transcriptional mechanisms and pathways that are operational during the onset of this condition. This provides a compendium of enhancer regions for instigating further mechanistic studies into the gene regulatory networks that are operational in this deadly disease.

## Acknowledgements

We thank Guanhua Yan for excellent technical assistance, and staff in the Bioinformatics and Genomic Technologies core facilities. We also thank Rebecca Fitzgerald, Cambridge on behalf of the OCCAMS network for providing the patient RNA-seq data and matched Barrett’s and OAC samples for KAS-seq analysis. We are grateful to Chuan He for providing us with N_3_-kethoxal. This work was funded by grants from the Manchester Cancer Research Centre (MCRC), the MRC (MR/V010263/1) and the Wellcome Trust (102171/B/13/Z).

## Author contributions

IA, SHY, CWB, YL performed the experiments and data analysis in this study; ADS contributed to the inception, design and supervision of the project. All authors contributed to manuscript preparation and/or critically appraised manuscript drafts.

## Materials and methods

### Cell culture and treatments

OE19 cells were cultured in RPMI 1640 (ThermoFisher Scientific, 52400), HEK293T cells were cultured in DMEM (ThermoFisher Scientific, 11965084). All media was supplemented with 10% foetal bovine serum (ThermoFisher Scientific, 10270). Cell lines were cultured at 37°C, 5% CO_2_ in a humidified incubator. Lapatinib (Selleckchem, S1028) treatments were performed for 24 hours and at a 500 nM final concentration.

### Luciferase reporter assays

Regions containing *VEGFAe*, *ELF3e*, *GATA6e* or *PAX8e* were amplified from OE19 genomic DNA using primers containing 20 bp overlap regions with the multiple cloning site of the pGL3 Promoter vector (Promega, E1761) for luciferase assays (Supplementary Table S4). Final vectors were assembled using HiFi assembly (NEB, E5520S) according to the manufacturer’s instructions to create pGL3 plasmids containing *VEGFAe*, *ELF3e*, *GATA6e* or *PAX8e* enhancer regions (pAS5014-pAS5017). Enhancer vectors were transfected into OE19 cells using the Amaxa™ Nucleofector™ II (Lonza) with Cell Line NucleofectorTM Kit V (Lonza, VCA-1003) using program T-020, according to manufacturer’s instructions. To conduct luciferase assays, 250 ng of enhancer vector was co-transfected alongside 50 ng of pCH110 (Amersham). Enhancer activity was assessed using the Dual-Light™ Luciferase & β-Galactosidase Reporter System (ThermoFisher Scientific, T1003) according to the manufacturer’s instructions.

### STARR-seq vector production

An integrating STARR-seq vector was designed based on the pLenti-FKBP-delCasp9-Puro vector [Pang & Snyder, 2020] and the hSTARR-seq_ORI vector (Addgene, 99296; [Muerdter et al, 2018]). Briefly, the lentiviral machinery was amplified from the pLenti-FKBP-delCasp9-Puro vector to create two PCR products (primer pairs ADS6967/ADS6968 and ADS6969/6970). The STARR reporter machinery was amplified from the hSTARR-seq_ORI vector to create two PCR products using primer pairs ADS6963/ADS6964 and ADS6965/6976. The four PCR products were then assembled in a 1:1:1:1 ratio using HiFi assembly (NEB, E5520S) according to the manufacturer’s instructions to create the pLenti-STARR vector (pAS5018). PCR primers are listed in Supplementary Table S4.

### STARR-seq plasmid library generation

To maintain library complexity, 8 Omni-ATAC or CUT&Tag tagmentation reactions of 50,000 OE19 cells each were conducted per library, as previously described [Corces et al, 2017; Kaya-Okur et al, 2020] except Nextera sample barcodes were only introduced using the forward primer (Supplementary Table S4) and with an altered nuclear extraction step for CUT&Tag libraries. For the CUT&Tag nuclear extraction, OE19 cells were initially lysed in Nuclei EZ lysis buffer (Sigma-Aldrich, NUC-101) at 4°C for 10 mins followed by centrifugation at 500 g for 5 mins. The subsequent clean-up was performed in a buffer composed of 10 mM Tris-HCl pH 8.0, 10 mM NaCl and 0.2% NP40 followed by centrifugation at 1300 g for 5 mins. Nuclei were then lightly cross-linked in 0.1% formaldehyde for 2 mins followed by quenching with 75 mM glycine followed by centrifugation at 500 g for 5 mins. Cross-linked nuclei were resuspended in 20 mM HEPES pH 7.5, 150 mM NaCl and 0.5 M spermidine at a concentration of 4-8×10^3^/ μL (2-4×10^4^ total). For 2-4 ×10^4^ nuclei, 0.5 μg of primary and secondary antibody were used with 1 μL of pA-Tn5 (Epicypher, 15-1017). Antibodies used for CUT&Tag: anti-BRD4 (abcam, ab128874), anti-H3K27ac (abcam, ab4729) and anti-MED1 (AntibodyOnline, A98044/10UG). Subsequent CUT&Tag stages were as previously described (Kaya-Okur et al., 2020).

Omni-ATAC or CUT&Tag libraries assembled into the pLenti-STARR vector using HiFi assembly (NEB, E5520S), as previously described [Arnold et al, 2013] and according to manufacturer’s instructions. 25 assembly reactions per library were conducted to maintain library complexity. Assembly reactions were pooled and cleaned using Ampure XP beads (Beckman Coulter Agencourt, A63881) at a 1.8X ratio of beads to input. Assembled libraries were eluted into 10 µL nuclease-free H_2_O. MegaX DH10B T1R electrocompetent bacteria (20 µL; ThermoFisher Scientific, C640003) were transformed with 150 ng of library, with a total of 8 transformations performed per library to maintain complexity. Bacteria were transformed using the Ec1 setting on the BioRad MicroPulser (BioRad, 165-2100) in 1 cm cuvettes, ensuring time constants between 4.5-5.0 ms. Bacterial cells were recovered in 750 µL of pre-warmed SOC medium and incubated for 1 hr in a 250 RPM shaker at 37°C. Post-recovery, bacterial cells were pooled and incubated overnight in 4L of Luria broth containing 100 µg/mL carbenicillin (Sigma, C1613). Plasmid libraries were collected using the Plasmid Giga Kit (Qiagen, 12191). An aliquot of the plasmid library was sequenced on an Illumina HiSeq 4000 System to assess complexity (University of Manchester Genomic Technologies Core Facility).

### STARR-seq library transduction and screening

Lentiviral particles were generated as previously described [Tiscornia et al, 2006]. Briefly, 2×10^6^ HEK293 cells were transfected with 15 µg pLenti-STARR library, 10 µg psPAX2 (Addgene, 12260) and 5 μg pMD2.G (Addgene, 12259) using PolyFect (Qiagen, 301107). Media was collected at 48 and 72 hours post-transfection and viral particles were precipitated using PEG-it™ Solution (System Biosciences, LV810A-1). To transduce, 1×10^7^ OE19 cells were treated with virus (MOI 0.7) and 5 μg/mL Polybrene (EMD Millipore, TR-1003). Polyclonal cells were selected for 2 weeks in 500 ng/μL puromycin (Sigma, P7255). After selection and growth to 5×10^7^ cells, 50% of cells were processed for gDNA isolation using the DNEasy Blood and Tissue Kit (Qiagen, 69504) and 50% processed for RNA isolation using the RNEasy Midi Kit with optional on-column DNAse digestion (Qiagen, 75144), as per manufacturer’s instructions. Sequencing-ready gDNA and RNA libraries were generated as previously described [Wang et al, 2018]. Briefly, polyA+ mRNA was isolated using the Oligotex mRNA Midi kit (Qiagen, 70042), as per manufacturer’s instructions. RNA was reverse-transcribed using Superscript III Reverse Transcriptase (ThermoFisher Scientific, 18080085) with 4 reactions per library (ensuring <4 μg per reaction) using a specific primer and according to manufacturer’s instructions (Supplementary Table S4). cDNA was pooled and cleaned up using Ampure XP beads (Beckman Coulter Agencourt, A63881) at a 1.8X ratio of beads to input, eluting in 80 μL nuclease-free H_2_O. cDNA and gDNA libraries were amplified by PCR using the NEBNext 2X Master Mix with the remaining sample barcode introduced using the reverse primer (Supplementary Table S4). 4 reactions were performed per library. Libraries were pooled and sequenced on an Illumina HiSeq 4000 System (University of Manchester Genomic Technologies Core Facility).

### STARR-seq data analysis

Initial STARR-seq data processing was performed similarly to ATAC-seq data processing previously described [Britton et al., 2017]. Briefly, reads were mapped to GRCh38 (hg38) using Bowtie2 v2.3.0 [Langmead & Salzberg, 2012] with the options: -X 2000 -dovetail. Mapped reads (>q30) were retained using SAMtools [Li et al., 2009]. Reads mapping to blacklisted regions were removed using BEDtools [Quinlan & Hall, 2010]. Peaks were called using MACS2 v2.1.1 [Zhang et al., 2008] with the following parameters: -q 0.01, -nomodel -shift -75 -extsize 150 -B –SPMR. A union peakset was formed from all plasmid library samples, using HOMER v4.9 mergePeaks.pl -d 250 [Heinz et al., 2010] as described previously [Rogerson et al., 2019]. STARR signal was ranked using featureCounts [Liao et al., 2014] by taking the sum of RNA libraries at regions above the DESeq2-defined count threshold and calculating signal over plasmid (ATAC-STARR-seq) or gDNA (CUT&Tag-STARR-seq). The changepoint package in R v3.6.0 was used to determine RNA and plasmid ranks, as well as the top-ranked regions for subsequent analysis. Differentially active STARR regions were subsequently determined from top-ranked regions using DESeq2 [Love et al., 2014]. A log_2_-fold change of ±0.3 and *P*-value_adj_ < 0.1 defined differential expression.

HOMER v4.9 was used for de novo transcription factor motif enrichment analysis. STARR+ regions were annotated to genes by the nearest gene model and genomic distribution profiled using HOMER v4.9 annotatePeaks.pl. A custom peakset of high-confidence STARR+ intersect regions was generated as described previously [Rogerson et al, 2019]. Differentially accessible/H3K27ac-marked high-confidence STARR+ intersect regions upon lapatinib treatment were determined using DESeq2 [Love et al., 2014]. A log_2_-fold change of ±0.2, and *P*-value_adj_ < 0.075 and 0.05 defined differential accessible and H3K27ac-marked regions, respectively.

### CUT&Tag processing and data analysis

OE19 cells were treated with 500 nM lapatinib or DMSO. After 24 hours, CUT&Tag library generation was performed as described above using an anti-H3K27ac antibody (abcam, ab4729). CUT&Tag libraries were pooled and sequenced on an Illumina HiSeq 4000 System (University of Manchester Genomic Technologies Core Facility). CUT&Tag data processing was performed as for ATAC-seq analysis described above. A union peakset was generated using HOMER v4.9 mergePeaks.pl -d 250 [Heinz et al., 2010] as described previously [Rogerson et al., 2019] and biological replicates were assessed for concordance (*r* > 0.80).

### KAS-seq processing and data analysis

24 hr-lapatinib or DMSO treated OE19 cells/cells from OAC tissue were prepared for KAS-seq. OE19 KAS-seq library generation was performed as described previously for bulk low input KAS-seq [Wu et al., 2020] except for altered nuclear extraction and labelling reactions, and using home-made Tn5 transposase as described previously [Picelli et al, 2014]. Nuclei were extracted and washed as described for CUT&Tag. Nuclei were then resuspended in nuclease-free H_2_O at a concentration of 1×10^4^/ μL. (2×10^5^ total) Labelling reactions were carried out in DNA LoBind® tubes (Eppendorf, 0030108051) using 5 mM N3-kethoxal (a gift from Chuan He) in PBS to a final volume of 50 μL for 15 mins at 37°C with 1000 RPM mixing in a thermomixer. Labelled gDNA was isolated using the PureLink™ Genomic DNA Mini kit (ThermoFisher Scientific, K182001) and eluted twice with 21.5 μL 25 mM K_3_BO_3_ pH 7.0. Purified DNA underwent tagmentation reaction with 2 µM of Tn5 in 1x TD buffer (33 mM Tris-OAC pH 7.8, 66 mM KOAc, 10 mM Mg(OAc)_2_ and 16% dimethylformamide) in a 25 µL reaction volume at 37°C for 30 mins. Subsequent enrichment was performed with 5 µL of Dynabeads Streptavidin C1 (ThermoFisher Scientific, 65001) and resuspended in 19.5 µL H_2_O.

Library amplification was performed by PCR with 20 µL beads, 0.5 µM i5 and i7 Illumina index primers (Illumina, 20027213) and NEBNext Ultra II Q5 Master Mix (NEB, M0544S) in a 50 µL final reaction volume. The PCR reaction was carried out at 72°C, 5 min; 95°C 10 min, 15 cycles at 98°C 10s, 63°C 30s, 72°C 60s and a final 72°C step for 2min. The final libraries were cleaned up using Zymo DNA Clean & Concentrator kit 5 (Zymo Research, D4014). KAS-seq libraries were pooled and sequenced on an Illumina HiSeq 4000 System (University of Manchester Genomic Technologies Core Facility). Three biological replicates were sequenced and checked for concordance (r > 0.80). KAS-seq data processing was performed as described previously [Wu et al., 2020], but with the MACS2 v2.1.1 --broad peak calling option.

For OAC and Barrett’s tissue KAS-seq, fresh frozen OAC 3 mm biopsies were collected at Cambridge University Hospitals NHS Trust (Addenbrooke’s Hospital) from patients undergoing endoscopy. The study was approved by the Institutional Ethics Committees and all patients gave individual informed consent. Tissue nuclei were extracted as previously described [Britton et al, 2017] and treated as described for OE19 cells.

KAS-seq data processing was performed as described previously [Wu et al., 2020]. Briefly, reads were mapped to the human genome GRCh38 (hg38) using Bowtie2 v2.3.0 [Langmead & Salzberg, 2012]. Mapped reads (>q30) were retained using SAMtools [Li et al., 2009]. Reads mapping to blacklisted regions were removed using BEDtools [Quinlan & Hall, 2010]. Peaks were called using MACS2 v2.1.1 [Zhang et al., 2008] with the following parameters: -q 0.01, -nomodel -shift -75 -extsize 150 -B –SPMR --broad. A union peakset was generated as described previously [Rogerson et al., 2019] and biological replicates were assessed for concordance (*r* > 0.80).

### Bioinformatics

Genome browser data was visualised using the UCSC Genome Browser [Kent et al., 2002]. Heatmaps and tag density plots of epigenomic data were generated the using deepTools [Ramirez et al., 2016] computeMatrix, plotProfile, plotCorrelation and plotHeatmap functions. Metascape [Zhou et al., 2019] was used for gene and disease ontology analysis of gene sets. The eulerr package in R v3.6.0 was used for generating Venn diagrams.

### Datasets

All data was obtained from ArrayExpress, unless stated otherwise and are listed in Supplementary Table S5. OAC tissue total RNA-seq data was obtained from: OCCAMS consortium (European Genome-Phenome Archive, EGAD00001007496)(Jammula et al., 2020). Lapatinib-treated OE19 ATAC-seq, lapatinib-treated OE19 RNA-seq and OE19 H3K27ac ChIP-seq was obtained from: E-MTAB-10334, E-MTAB-10304 and E-MTAB-10302, respectively [Ogden et al., 2022]. OE19 KAS-seq and CUT&Tag data was obtained from E-MTAB-11357 and E-MTAB-11356, respectively [Ahmed et al, 2023]. The 4,600 test eRNA set was obtained from Ahmed et al, 2023.

### Data access

STARR-seq data (E-MTAB-14083 and E-MTAB-14155), OAC and Barrett’s tissue KAS-seq (E-MTAB-14091), and KAS-seq data in OE19 cells (E-MTAB-14095 and E-MTAB-14098) are deposited in ArrayExpress and listed in Supplementary Table S5.

---

**Figure S1.**
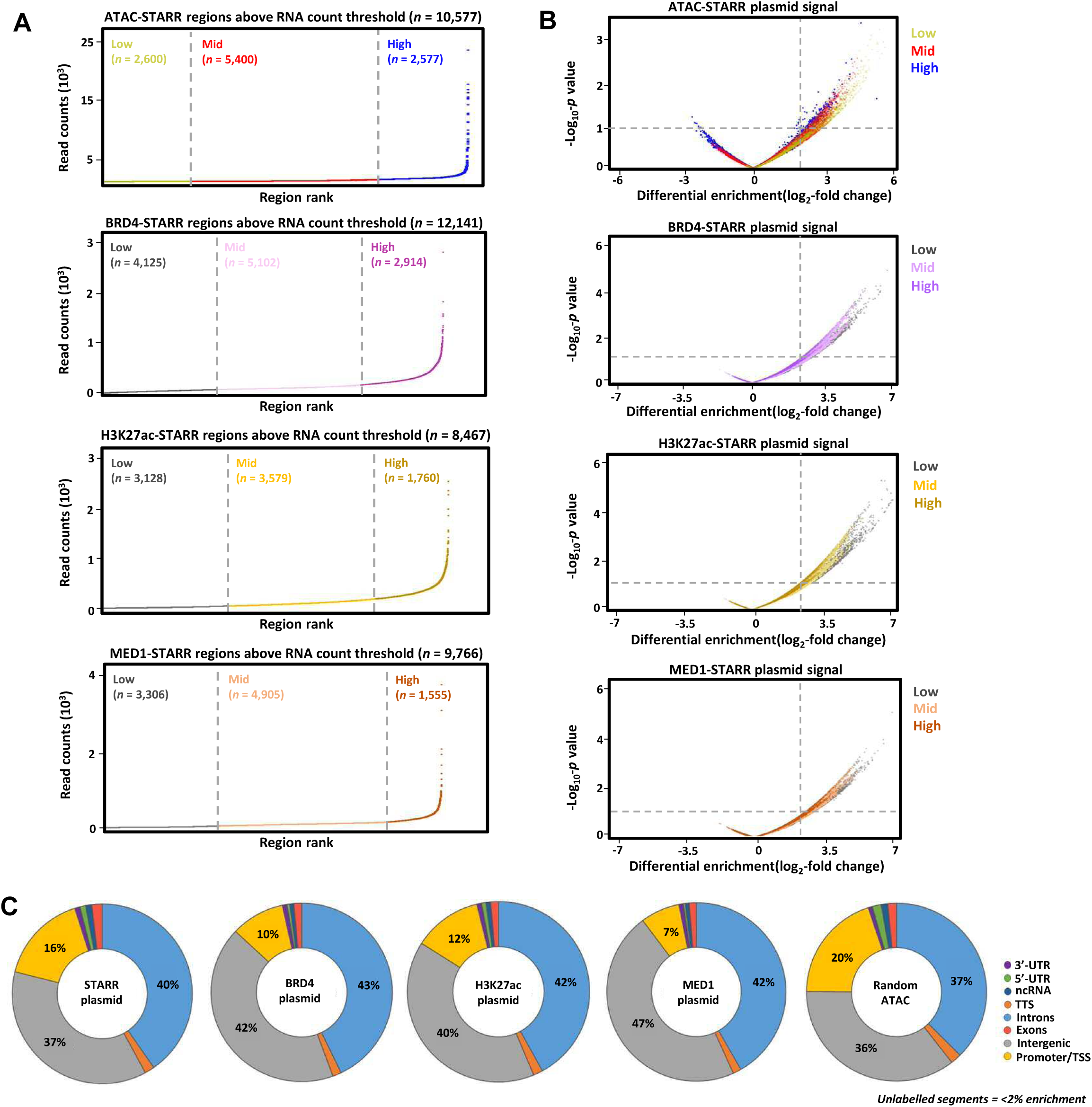
STARR-seq identifies regions of the genome capable of driving self-transcription. (A) Plots displaying ranked RNA count signal for ATAC, BRD4, H3K27ac and MED1-STARR-seq assays for detectable RNA-producing regions. Regions have been trisected into low, mid and high RNA count regions. (B) Volcano plots displaying the RNA signal for ATAC, BRD4, H3K27ac and MED1-STARR-seq assays with regions colour coded by plasmid signal strength. (C) Genomic distribution of the STARR plasmid library for ATAC, BRD4, H3K27ac and MED1-STARR-seq assays, and random ATAC regions (n=10,000).

**Figure S2.**
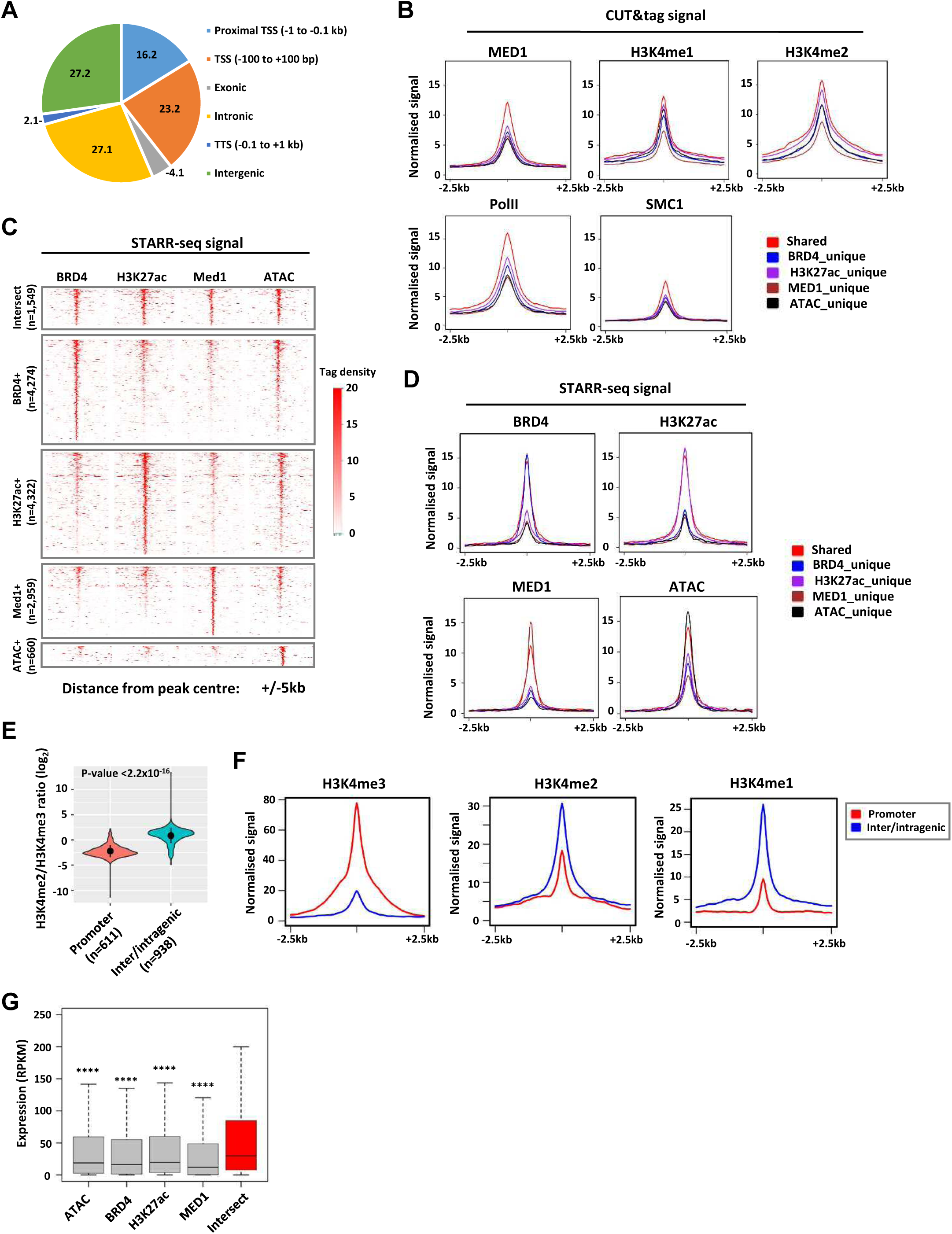
High confidence STARR+ regions are associated with features of active regulatory elements. (A) Genomic distribution of the 1,549 high confidence STARR+ regions. (B and D) Metaplots of CUT&Tag signals (B) and STARR-seq signals (D) for the indicated factors/marks in 1,549 STARR+ intersect regions compared to STARR+ unique regions in ATAC, BRD4, H3K27ac and MED1-STARR-seq assays. (C) Heatmaps showing STARR-seq signal in OE19 cells at the 1,549 STARR+ intersect regions compared to STARR+ unique regions in ATAC, BRD4, H3K27ac and MED1-STARR-seq assays. (E) Violin plots displaying the ratio of H3K4me2:H3K4me3 ratios at intergenic regions and promoters (−1kb to +0.1kb) within the 1,549 STARR+ intersect regions (*p*-value is shown; Student’s t-test). (F) Metaplots of OE19 cell H3K4me3, H3K4me1 and H3K4me3 CUT&Tag signal at promoter proximal (−1kb to +0.1kb; n=611) and distal regulatory regions (n=938) within the 1,549 STARR+ intersect regions. (G) Box plots comparing the expression of genes in OE19 RNA-seq samples annotated to unique BRD4, H3K27ac, MED1-STARR+ regions against genes annotated to the 1,549 STARR+ intersect regions (*p*-value is shown; ****=<1×10^−5^, t-test).

**Figure S3.**
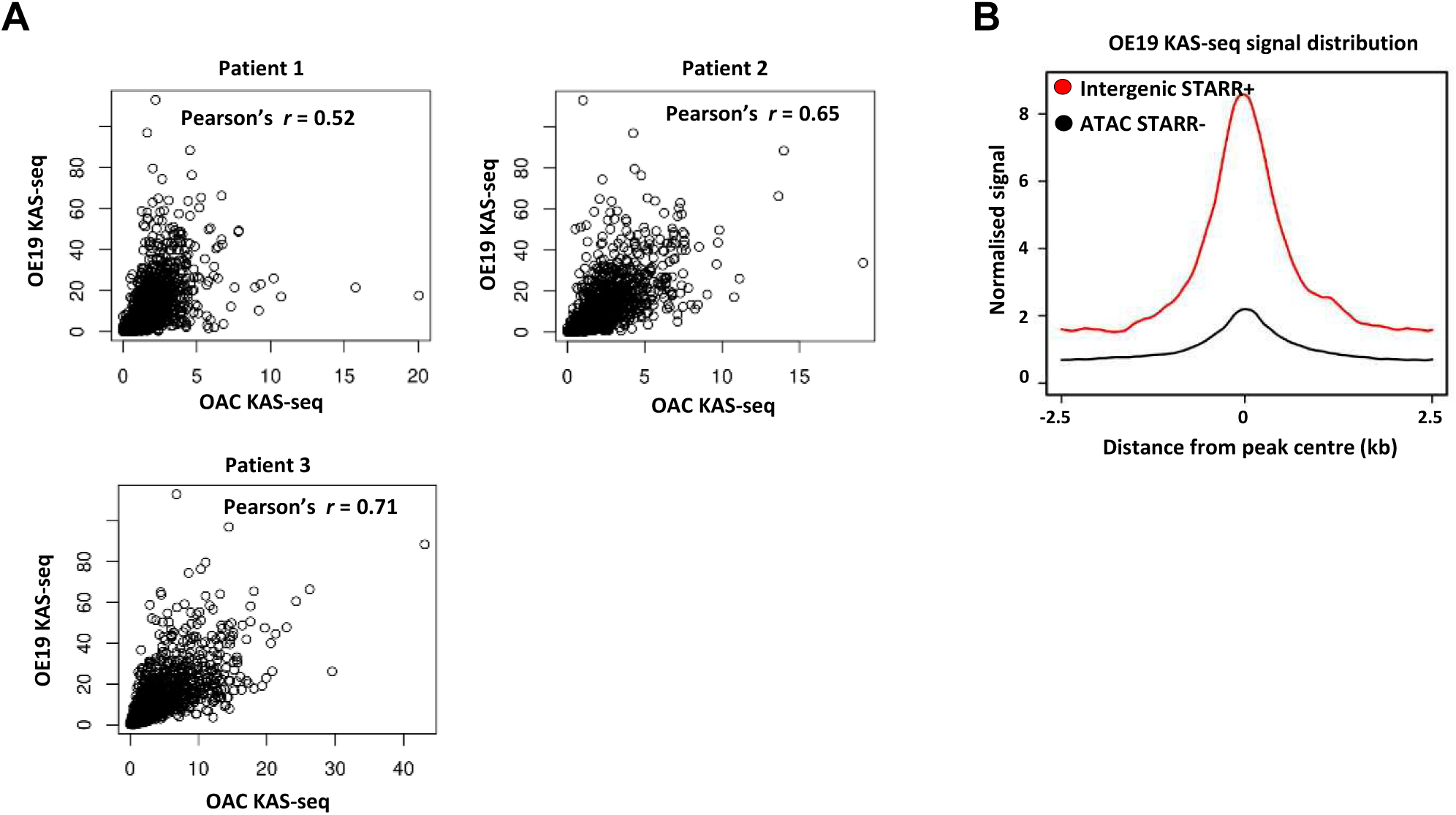
High confidence STARR+ regions are associated with transcription-associated DNA opening. (A) Correlation plot of KAS-seq signal for OE19 cells and OAC tissue from three different patients (Pearson’s correlation coefficient is shown). (B) Metaplot of OE19 cell KAS-seq signal at the 566 STARR+ intersect regions found in intergenic regions compared to 2,303 randomly selected STARR-ATAC peaks from OE19 cells.

**Figure S4.**
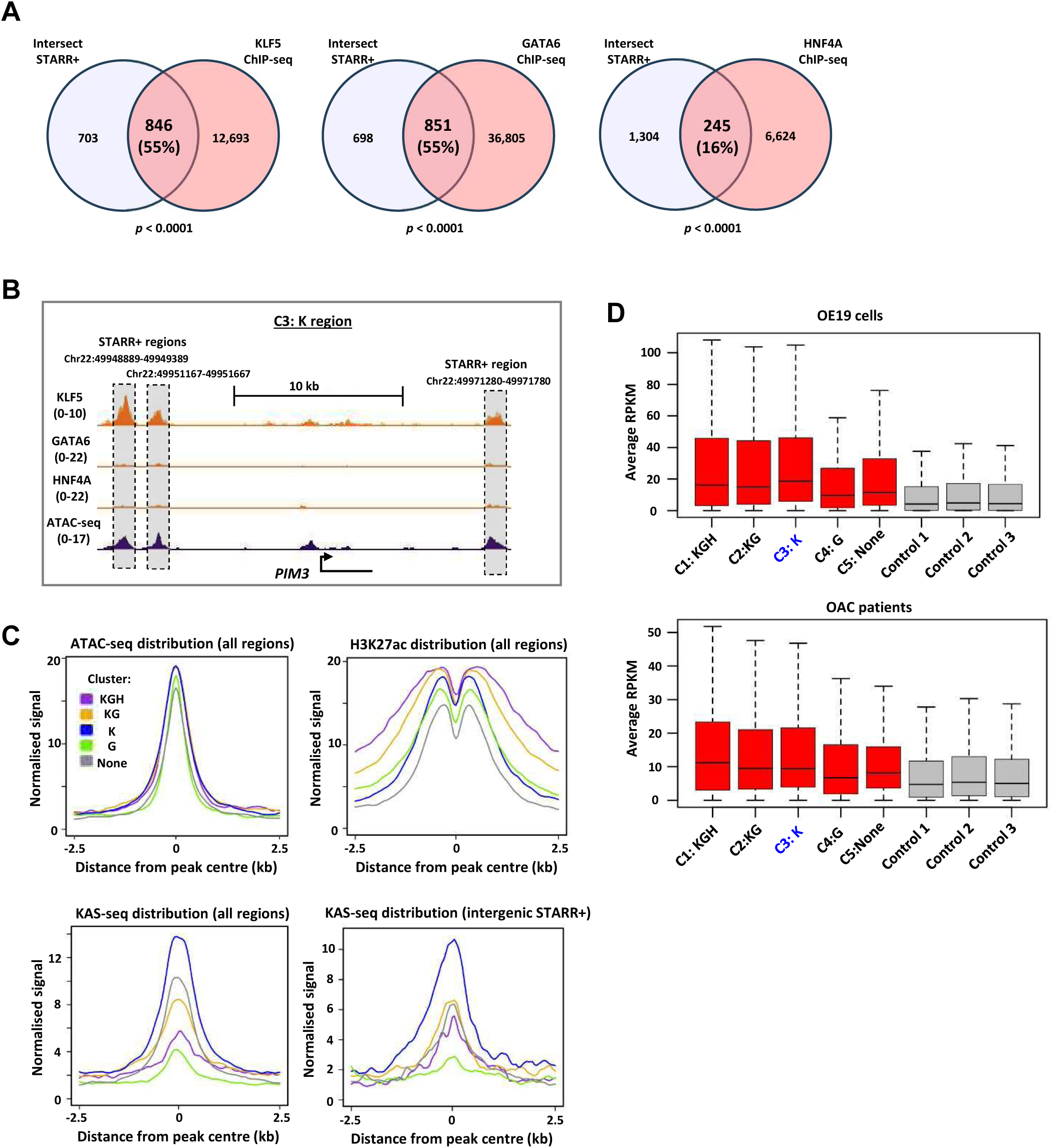
High confidence STARR+ regions are enriched for binding to core OAC transcription factors and active chromatin features. (A) Venn diagrams of the overlap in ChIP-seq peaks for the transcription factors KLF5, GATA6 and HNF4A in OE19 cells and the 1,549 high confidence STARR+ regions. (B) Genome browser view of KLF5, GATA6 and HNF4A ChIP-seq, and ATAC-seq signal at C3:K cluster STARR+ intersect regions (highlighted) surrounding the *PIM3* locus. (C) Average tag density plots of ATAC-seq, H3K27ac ChIP-seq or KAS-seq signal in OE19 cells for each of the 5 clusters in either all of the high confidence STARR+ regions or in intergenic regions only (bottom right). (D) Boxplot of average expression values of the genes nearest to the 1,549 STARR+ intersect regions from clusters C1-5 or 3 control sets of randomly selected ATAC-seq peaks, in OE19 cells (top) or 28 OAC patient samples (bottom). All red bars are significantly higher than the control grey bars; P-values all <0.001 in OE19 cells and <0.05 in patient samples.

**Figure S5.**
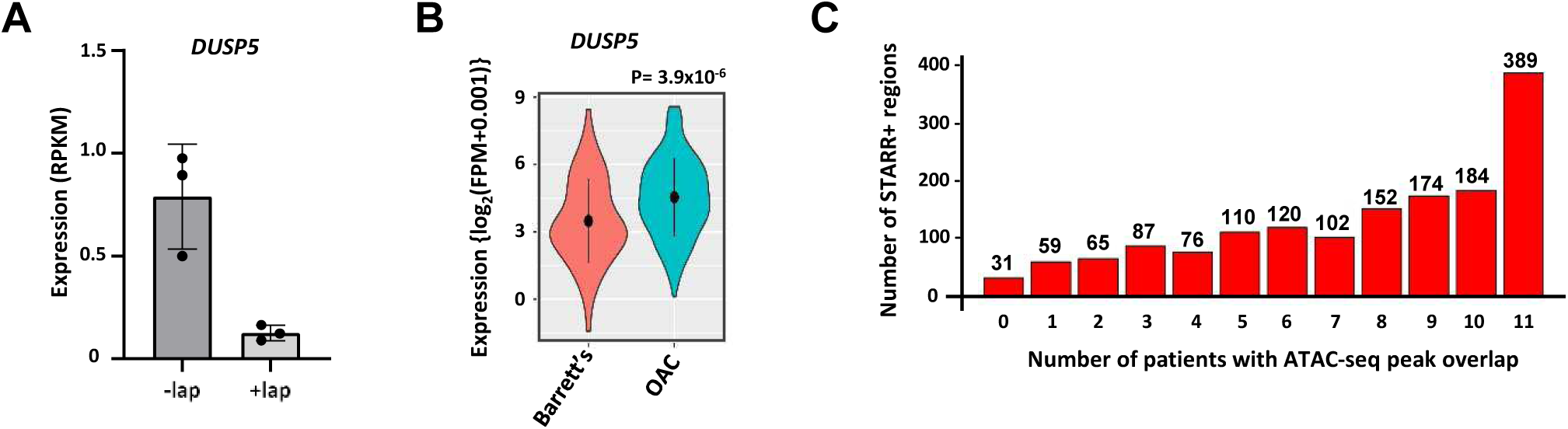
A subset of high confidence STARR+ regions are responsive to treatment with the ERBB2 inhibitor lapatinib. (A) *DUSP5* expression from OE19 RNA-seq data (Ogden et al., 2022) in the presence or absence of lapatinib treatment for 24 hrs (n=3). (B) Violin plot of *DUSP5* expression in RNA-seq data from OAC and Barrett’s patients. (C) Numbers of STARR+ regions overlapping with open chromatin peaks in 11 OAC patient ATAC-seq datasets. Data show the number of STARR+ regions (y-axis) versus the number of patients containing an overlap (x-axis).

